# Humanized CD36 mouse model supports the preclinical evaluation of therapeutic candidates targeting CD36

**DOI:** 10.1101/2023.01.09.523279

**Authors:** Xiulong Xie, Zhenlan Niu, Linlin Wang, Xiaofei Zhou, Xingyan Yu, Hongyan Jing, Yi Yang

**Author notes:** These authors contributed equally to this work (Xiulong Xie, Zhenlan Niu, Linlin Wang).

## Abstract

CD36 (also known as scavenger receptor B2) is a multifunctional receptor that mediates lipid uptake, advanced oxidation protein products, and immunological recognition, and has roles in lipid accumulation, apoptosis, as well as in metastatic colonization in cancer. CD36 is involved in tumor immunity, metastatic invasion, and therapy resistance through various molecular mechanisms. Targeting CD36 has emerged as an effective strategy for tumor immunotherapy. In this study, we have successfully generated a novel CD36 humanized mouse strain where the sequences encoding the extracellular domains of the mouse *Cd36* gene were replaced with the corresponding human sequences. The results showed that CD36 humanized mice expressed only human CD36, and the proportion of each lymphocyte was not significantly changed compared with wild-type mice. Furthermore, CD36 monoclonal antibody could significantly inhibit tumor growth after treatment. Therefore, the CD36 humanized mice provided a validated preclinical mouse model for the evaluation of tumor immunotherapy targeting CD36.

## Introduction

The transmembrane protein CD36 (also known as scavenger receptor B2) is a membrane glycoprotein expressed on the cell surface in multiple cell types, including platelets, mononuclear phagocytes, adipocytes, hepatocytes, myocytes, and some epithelia[1]. Clinical studies have found significant upregulation of CD36 expression in tumor tissues of cervical cancer[2, 3], gastric cancer[4], hepatocellular carcinoma[5], and ovarian cancer[6], which promotes tumor growth, metastatic invasion, and therapy resistance of these tumors through CD36-mediated lipid metabolism.[7]. In addition, the tumor microenvironment (TME) exerts several types of metabolic stress on the infiltrating immune cells, including acidosis, hypoxia, and nutritional deficiencies[8]. Upregulation of CD36 expression not only helps tumor-infiltrating Treg cells adapt to TME and further exert their immunosuppressive effects[9], but also causes tumor-infiltrating CD8^+^ T cells to induce lipid peroxidation and iron death due to lipid accumulation, thus affecting the anti-tumor ability of CD8^+^ T cells[10, 11].

Colorectal cancer (CRC) is the third most common cause of cancer-related death worldwide, and the malignant transformation of benign adenomas and polyps is the cause of CRC[12]. CD36 expression was significantly higher in primary CRC tumor tissue than in normal colonic mucosa, and 5-year survival was lower in CRC patients with high CD36 mRNA expression than in those with low CD36 mRNA expression [13].CD36 is involved in CRC development by promoting proteasome-dependent ubiquitination of GPC4 to inhibit the β-catenin/c-myc axis[14]. The non-coding RNA (lncRNA) TINCR was found to inhibit miR-107 expression and activate CD36, which further inhibited CRC cell proliferation and promoted CRC cell apoptosis[15]. Inhibition of FASN was found to upregulate CD36 expression in fatty acid synthase (FASN) knockout CRC cells and CRC models based on transgenic mice with hetero- and homozygous deletions of FASN, thereby promoting the proliferation of CRC cells[13]. CRC with high metastatic potential expresses higher levels of CD36, which promotes CRC metastasis by upregulating MMP28 and increasing E-calmodulin cleavage[16]. Lipid droplets (LD) are a common feature of cancer cell adaptation to TME acidosis and a key driver of increased cancer cell aggressiveness[17]. It was found that acidosis of TME induces plasma membrane transport of CD36 via TGF-β2, which promotes LD formation and enhances metastasis and invasion of CRC[18]. Therefore, CD36 is closely associated with CRC development, growth, tumor immunity, and metastatic invasion, suggesting that inhibition of CD36 may be necessary to improve the efficacy of FASN-targeted therapy.

Humanized mice include immunodeficient mice xenografted with human cells or tissues as well as mice expressing human gene products[19]. PDX/CDX models based on immune reconstituted mice or target gene humanized mice are the most used for preclinical *in vivo* efficacy evaluation of antibodies for tumor immunotherapy. However, immune reconstituted mice have limitations such as species specificity of histocompatibility complex (MHC) antigens, underdevelopment of the immune system, impaired class switching and affinity maturation of immunoglobulins[20]. Therefore, a murine model used to conduct preclinical testing of anti-hCD36 Abs would be an invaluable tool for defining their mechanism of action and potential clinical utility. In this study, we described the generation and characterization of the humanized CD36(hCD36) mouse strain and validation of their use in studying CD36-targeting therapies for potential application in *in-vivo* anti-tumor activity.

## Materials and methods

### Generation of humanized CD36 mice

Based on C57BL/6 genetic background mice, exons 4 to part of 15 encodings the extracellular region of murine *Cd36* were replaced with exons 3 to 14 of human *CD36*.

### Phenotypic analyses of humanized CD36 mice

#### Analysis of human CD36 mRNA expression

Lung tissues of homozygous hCD36 mice (Unless otherwise stated, hCD36 mice below refer to homozygous hCD36 mice) and wild-type (WT) C57BL/6J were harvested for extracting total RNA using RNAprep Pure Cell / Bacteria Kit (TIANGEN, DP430). The mRNA expression of hCD36 and mCD36 was determined using GAPDH as the internal control. Primer annealing temperatures and number of cycling were set as follows: initial 94 °C for 2 min, followed the first step by 15 cycles of 98 °C for 10 s, 67 °C for 30 s, and 68 °C for 30 s, the second step by 25 cycles of 98 °C for 10 s, 57 °C for 30 s, and 68 °C for 30 s; an additional extension at 68 °C for 5 min; and finally held at 16 °C. The primer sequence of RT-PCR was designed as follows Table 1.

**Table 1.**
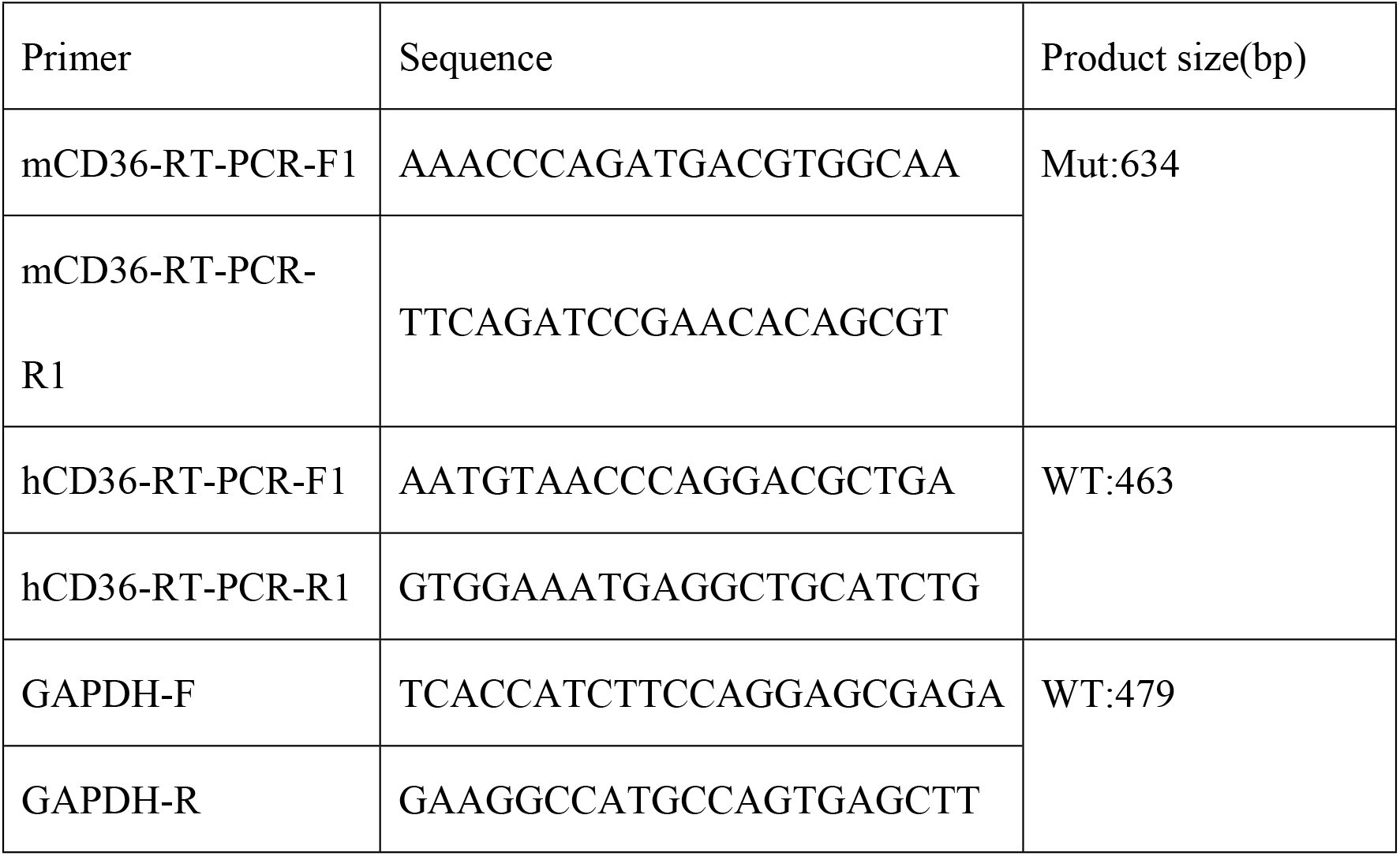
The primer sequence of RT-PCR.

#### Protein expression analysis of CD36 and detection of immunophenotyping

CD36 expression was examined in bone marrow derived from WT mice and heterozygous hCD36 mice, and hCD36 mice (peritoneal exudative macrophages) using Anti-mouse CD36 Antibody-APC (Biolegend, 102611) and Anti-mouse CD36-PE (eBioscience^TM^, 17-0362-82). And hCD36 was detected with anti-human CD36 antibody (Biolegend, 336205) in peritoneal exudative macrophages.

We next analyzed the expression of CD36 on immune cells in the blood of WT mice and hCD36 mice using flow cytometry. ACK lysis buffer (Beyotime, China) was first added to the anticoagulated blood to remove red blood cells. These cells were then co-incubated with a mixture of LD-NIR (Biolegend, USA) and anti-mCD16/32 (Biolegend, USA, clone 93) respectively for 10 min at 4°C for deadwood staining and blocking of non-specific binding. Finally, immune cells in these cell suspensions were stained at 4°C. After each staining, cells were washed with PBS to remove unbound labeled antibodies. After staining, multicolor flow cytometry of the cells was performed using Attune NxT (Thermo Fisher Scientific, USA) and analysis of the data was performed using FlowJo 10.

Finally, we analyzed the development, differentiation, and distribution of immune cells in the spleen, lymph nodes, and blood of WT mice and hCD36 mice, respectively. The spleen cell suspensions and lymph node cell suspensions were processed and analyzed with the same procedure as anticoagulated blood.

Listed below are the antibodies used for flow cytometric analysis of mouse cell-surface molecules: ① Purchased from Biolegend(USA): Anti-CD45-BV510(clone 30-F11);Anti-Ly-6G/Ly-6c(Gr-1)-PerCP(clone RB6-8C5);Anti-CD4-BV421(clone GK1.5);Anti-F4/80-FITC(clone BM8);Anti-CD8a-PE(clone 53-6.7); Anti-Foxp3-APC(clone FJK-16S); Anti-CD19-FITC(clone 6D5); Anti-TCR βchain-PerCP/Cy5.5 (clone H57-597); Anti-CD11c-BV605(clone N418); Anti-CD11b-PE (clone M1/70); Anti-CD11b-V450 Rat(clone V450); Purified Anti-CD16/32. Purchased from BD Pharmingen(USA): Anti-NK1.1-PE-Cy7 (clone PK136); Anti-TCR βchain-APC (clone H57-597). ② Purchased from eBioscience™: CD36 Monoclonal antibody-APC(clone HM36).

Listed below are the antibodies used for flow cytometric analysis of human cellsurface molecules: ①Purchased from Biolegend(USA): Anti-hCD36-PE(clone 5-271).

Other reagents: PE Mouse lgG2a, κ Isotype Ctrl(FC) Antibody (Biolegend, clone MOPC-173);Armenian Hamster lgG Isotype Control-APC(Invitrogen, clone eBio299Arm)

### Routine mouse blood test

Peripheral blood of female WT and hCD36 mice (n=8, 6-8 weeks old) was collected into EDTA blood collection tubes after anesthesia with sodium pentobarbital. The counts of red blood cells (RBC), white blood cells (WBC), neutrophils (NEUT#), lymphocytes (LYMPH#), monocytes (MONO#), hemoglobin (HGB), and platelets (PLT) in the peripheral blood of mice were determined using a fully automated modular blood fluid analyzer (Sysmes, XN-1000); the pressure of red blood cells (HCT), the mean red blood cell volume (MCV) and red blood cell distribution width (RDW); mean platelet volume (MPV), hemoglobin content (MCH) and mean hemoglobin concentration (MCHC).

### Biochemical examination of mouse peripheral blood

Peripheral blood of female WT and hCD36 mice (n=8, 6-8 weeks old) was collected into heparin collection tubes after anesthesia with sodium pentobarbital, and the supernatant was collected after centrifugation. The concentrations of alanine aminotransferase (ALT), aspartate aminotransferase (AST), alkaline phosphatase (ALP), albumin (ALB), glucose (GLU), urea (UREA), creatinine (CREA), total cholesterol (TC), triglycerides (TG), total protein (TP), alkaline phosphatase (ALP), serum amylase (AMY), phosphorus (P), creatine kinase (CK), high-density lipoprotein (HDL-C) and low-density lipoprotein (LDL-C) in the peripheral blood of mice were determined using a fully automatic biochemical analyzer (Hitachi, 3110).

### *In vivo* efficacy evaluation of anti-human CD36 antibody

Six-to eight-week-old humanized female hCD36 mice were housed in the specific-pathogen-free (SPF) barrier facility of the Animal Center of Biocytogen Pharmaceuticals (Beijing) Co., Ltd. in the individually ventilated cage. All experimental animal procedures were following the Institutional Animal Care and Use Committees (IACUC) guidance. Mice were euthanized with CO_2_. Mice were euthanized with CO_2_ to minimize or alleviate the animals’ suffering. 5E5 Murine colon cancer MC38 cells were subcutaneously implanted into hCD36 mice on the right dorsal side in a volume of 0.1ml per mouse. The tumor volume was measured once a day from day 0 after inoculation, and the tumor volume was calculated by the formula: 0.5 × long diameter × short diameter^2^. Mice were randomly grouped as tumors reached an average of 100 mm^3^. Then mice were treated with PBS, anti-hCD36 antibody (subsequently abbreviated as 1G04) through i.p. injection. Animal well-being and behaviors were monitored during the experiment process. We measured the tumor volume and weight of the animals twice weekly. The antitumor efficacy is expressed as tumor growth inhibition in terms of tumor volume (TGITV). The TGITV in percent was calculated as below: *TGI_TV_*(%) = [1 - (*T_i_*, - T_0_)/(*C_i_*, - C_0_)] × 100; Where *T_i_* = mean tumor volume of the drug-treated group on the final day of the study, T_0_= mean tumor volume of the drug-treated group on first dosing day, *C_i_* = mean tumor volume of the control group on the final day of the study, C_0_= mean tumor volume of the control group on the first dosing day.

### Statistical analyses

Mean±SEM was used to assess the results. Statistical analysis was performed using SPSS 19 and graphical plotting of data was performed using Graph Pad Prism 7 software. Student’s t-test and one-way analysis of variance (ANOVA) were used for the comparison of all data. Statistical significance was required to meet a *P* value < 0.05. \**p* <0.05, \*\**p* < 0.01, \*\*\**p* < 0.001, and \*\*\*\**p* < 0.0001.

## Results

### Generation of humanized CD36 mouse model and expression of human CD36 in humanized CD36 mice

The hCD36 mice established based on the C57BL/6 mouse background were obtained by replacing exons 4 to part of 15 of the extracellular region of CD36 encoding the mouse with human exons 3 to 14 (Fig 1A). We assumed that hCD36 mice only express hCD36, while heterozygous hCD36 mice express both hCD36 and mCD36. We detected CD36 expression in the bone marrow of heterozygous hCD36 mice and found that it expressed both mCD36 and hCD36, while WT mice only expressed mCD36 (results not shown). We analyzed the expression of CD36 mRNA in the lung tissue of hCD36 mice by RT-PCR. The results revealed that only hCD36 mRNA was detected in hCD36 mice compared to WT mice (Fig 1B). We then examined hCD36 protein expression in peritoneal exudative macrophages of hCD36 mice and found that only mCD36 was detected in WT mice, while only hCD36 was detected in hCD36 mice (Fig 1C).

**Fig 1.**
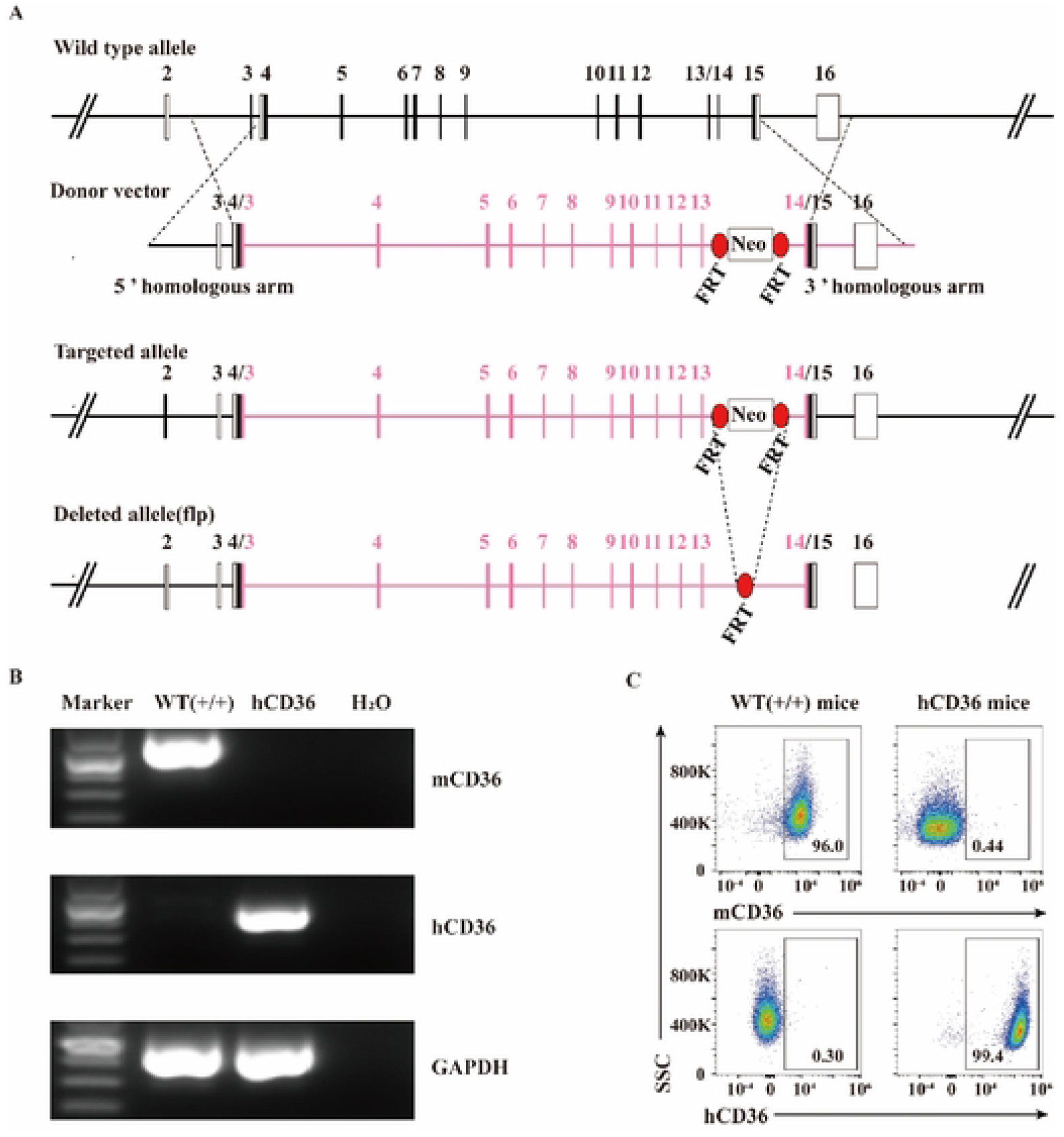
Characterization of the hCD36 mouse model. A: Generation of the hCD36 mouse strain. B: Expression of CD36 mRNA in hCD36 mice; C: Expression of CD36 on peritoneal exudative macrophages in hCD36 mice (n=1,7-week-old).

We further evaluated the expression pattern of hCD36 on different immune populations in the humanized mice and compared it with the expression of hCD36 in humans. Similar to the pattern found on human immune cells, hCD36 is expressed on macrophages, granulocytes, B cells, and T cell subsets in the hCD36 mice[21–23] (Table 2, S1 Fig). Altogether, our data demonstrate that the hCD36 mice recapitulate the expression pattern of the human CD36 on peripheral blood mononuclear cells (PBMCs), and validate the use of this mouse to study human CD36 receptor as the therapeutic target for immune therapy.

**Table 2.**
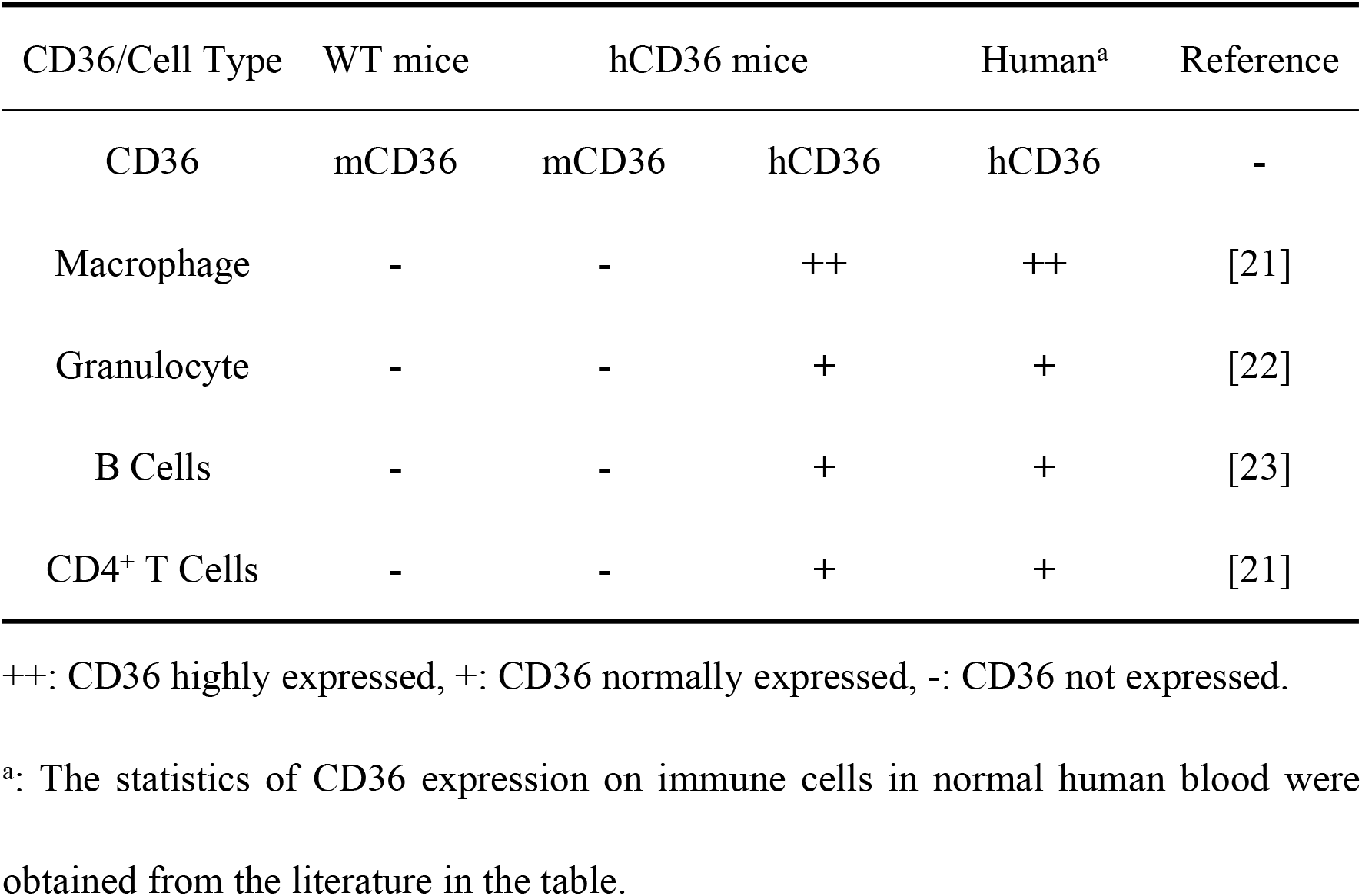
Expression of CD36 on immune cells in the blood of hCD36 mice.

### Analysis of leukocyte subpopulations and T cell subpopulations

To investigate whether the humanized hCD36 could affect the immune system of mice, we next analyzed the leukocyte subpopulation in the spleen of hCD36 mice by flow cytometry. As shown in Figs 2A and B, the development, differentiation, and distribution of leukocyte subpopulations such as T cells, B cells, natural killer cells (NK), monocytes, dendritic cells (DC), and mononuclear macrophages in the spleen of hCD36 mice were not statistically different from WT mice (p > 0.05). T cell subpopulations such as CD4^+^, CD8^+^ T cells, and Treg cells in the spleen of hCD36 mice were also similar to WT mice (p > 0.05) (Figs 2C and D). In addition, we further analyzed the development, differentiation, and distribution of leukocytes subpopulations and T cell subpopulations in the blood and lymph nodes of hCD36 mice. We found that leukocytes subpopulations and T cell subpopulations of blood and lymph nodes were not significantly changed as compared with WT mice(p > 0.05). (S2 and S3 Figs). Collectively, these results indicate that the development, differentiation, and distribution of immune cells in hCD36 mice are not impaired, and hCD36 mice possess normal immune functions.

**Fig 2.**
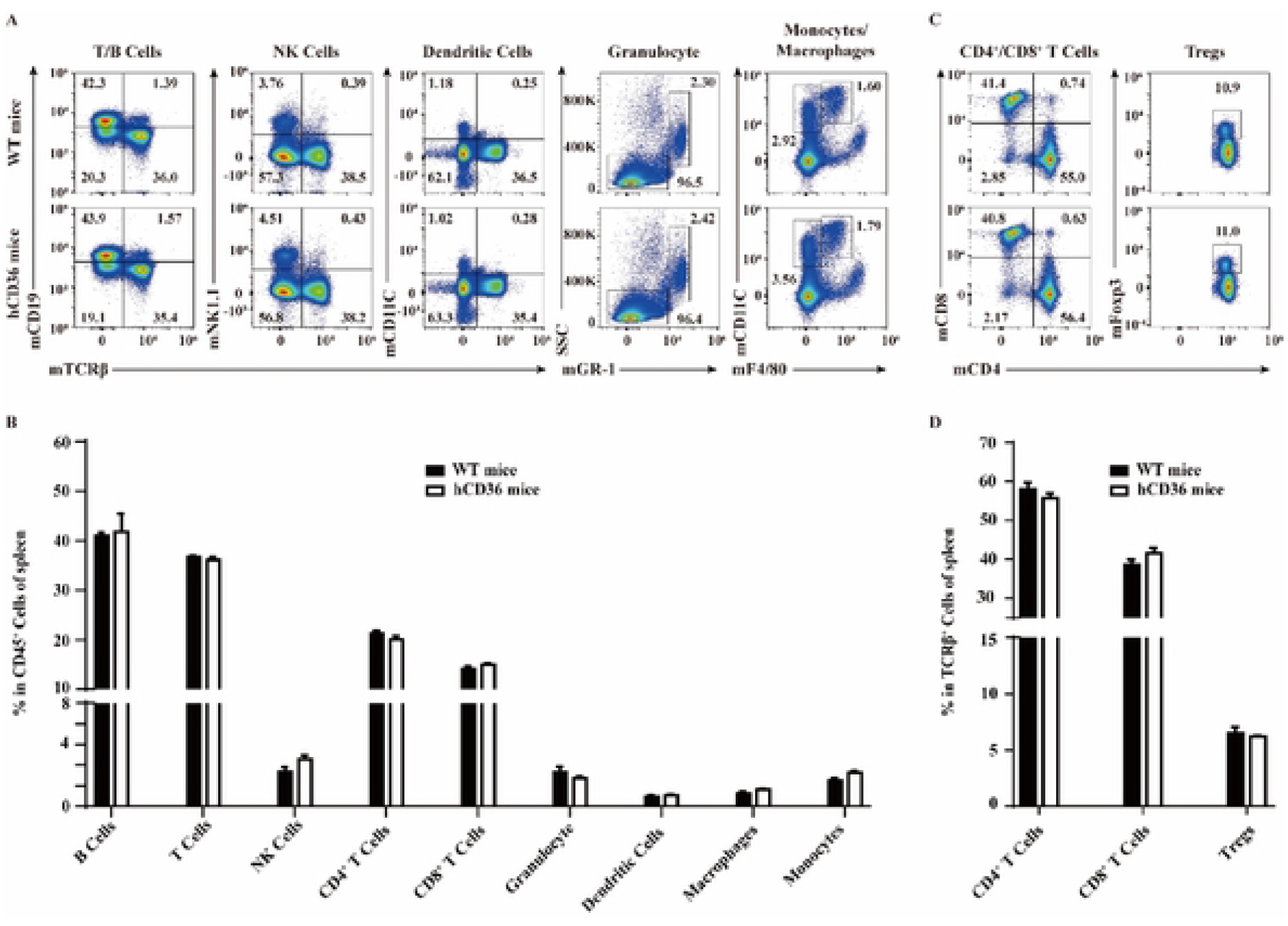
Analysis of immune cells subpopulation in the spleen of hCD36 and WT mice(n=3,8-week-old). A, B: The frequency of T cells, B cells, NK cells, Monocyte, DC cell, and macrophage cells in mCD45 cells. C, D: The frequency of CD4+ T cells, CD8+ T cells, and Treg cells in mCD3 cells.

### Analysis of routine blood and blood biochemistry

CD36 expression in non-immune cells mainly includes platelets, immature erythrocytes, podocytes, skeletal muscle cells, adipocytes, and cardiomyocytes[24]. We next examined whether blood cell composition and morphology were affected by CD36 humanization. As shown in Fig 3A, hCD36 mice were measured similarly to WT mice, indicating that humanization does not alter hematocrit composition and morphology.

**Fig 3.**
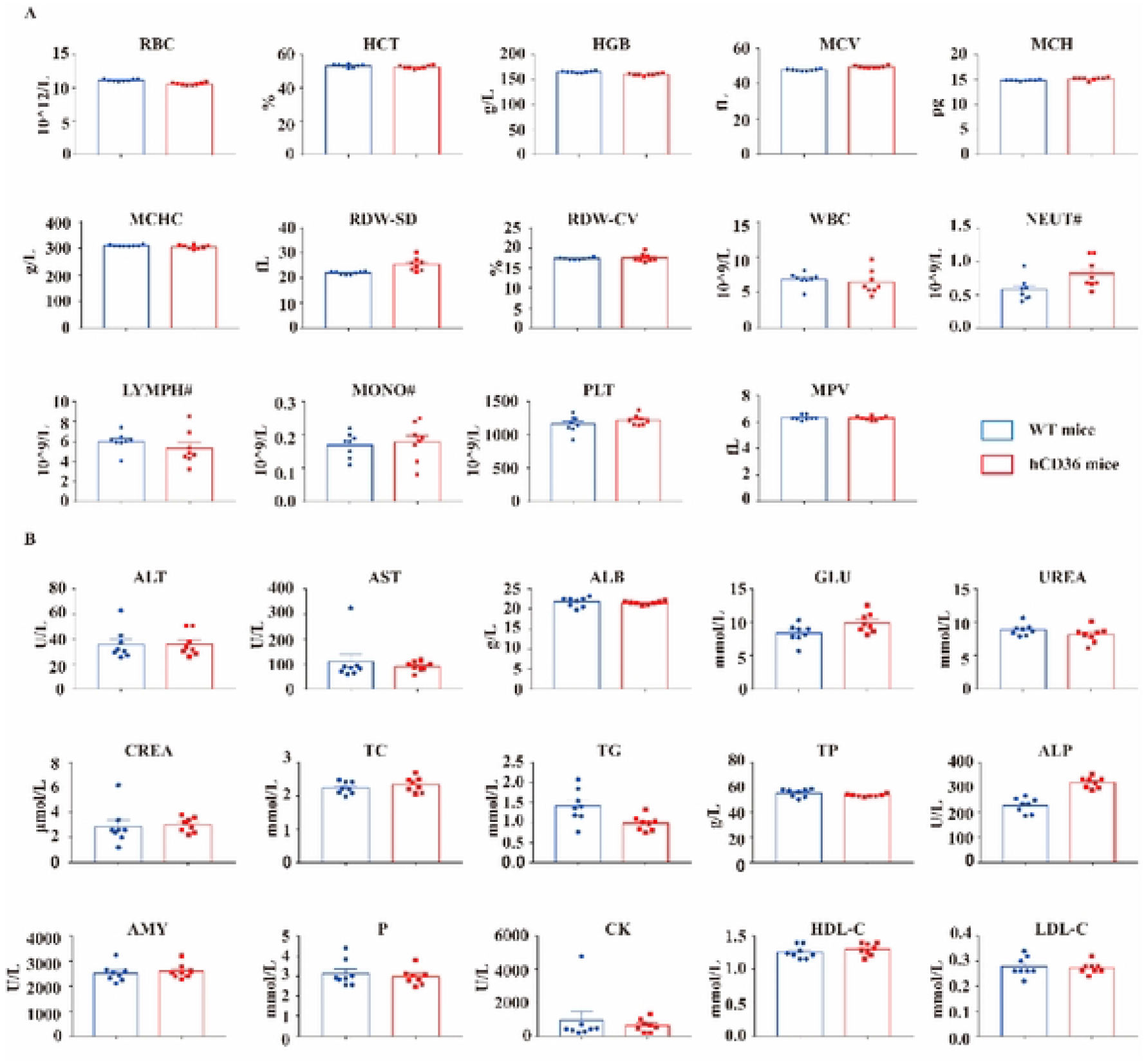
Analysis of complete blood count and blood biochemical in hCD36 and WT mice (n=3,6~8-week-old). A: Analysis of complete blood count in mice; B: Analysis of blood biochemistry in mice.

Notably, cardiomyocyte-specific CD36 knockout mice exhibited a significant reduction in cardiac FA uptake and intramyocardial TG content[25]. Hepatocyte-specific CD36 knockout mice exhibited high-fat diet-induced hepatic steatosis and diminished insulin resistance, and blood biochemical assays suggested a progressive decrease in AST and ALT[26]. CD36 also contributes to the progression of chronic kidney disease by mediating renal lipid deposition, lipid peroxidation, and endocytosis of multiple substances by renal cells[27]. Endothelial cell-specific CD36 knockout mice also exhibit elevated triglyceride levels, reduced total cholesterol, and increased glucose clearance[28]. These studies suggest that CD36 is closely associated with the normal function and disease development of organs such as the heart, liver, and kidney in mice. We next evaluated the effect of CD36 humanization on the normal function of these organs in hCD36 mice. There was no difference in biochemistry parameters between hCD36 mice and WT mice, indicating that humanization does not alter the health of organs such as the heart, liver, and kidney (Fig 3B).

### *In vivo* efficacy evaluation of anti-human CD36 antibody

The positive drug 1G04 is a human-mouse chimeric antibody, which was shown to slow down tumor growth and reduce the metastatic area of colon cancer in MC38 tumor-bearing WT mice [29]. To assess the cross-reactive ability of 1G04, we examined the binding of 1G04 to mCD36/hCD36 *in vitro* by flow cytometry, and the results were as expected that the 1G04 demonstrated detectable binding to both mCD36 and hCD36 (Fig 4B). To demonstrate that the enhancement of anti-tumor immunity could be modulated by therapeutic blockade of CD36, hCD36 mice with established MC38 wild-type tumors were treated with 1G04 separately at 3mg/kg or 10mg/kg. 1G04 treatments at dosage of 10mg/kg and 3mg/ kg achieved significant tumor growth inhibition (TGI) at 47.3% and 31.6%, respectively compared with the control group(Fig 4C). And neither the low-dose group nor the high-dose group induced significant body weight change in the experimental animals compared with the control group(p > 0.05)(Fig 4D). Therefore, we successfully validated the inhibitory effect of 1G04 on colon cancer tumor growth in hCD36 mice. The hCD36 mice could be used for preclinical *in vivo* efficacy evaluation of Anti-hCD36 antibody.

**Fig 4.**
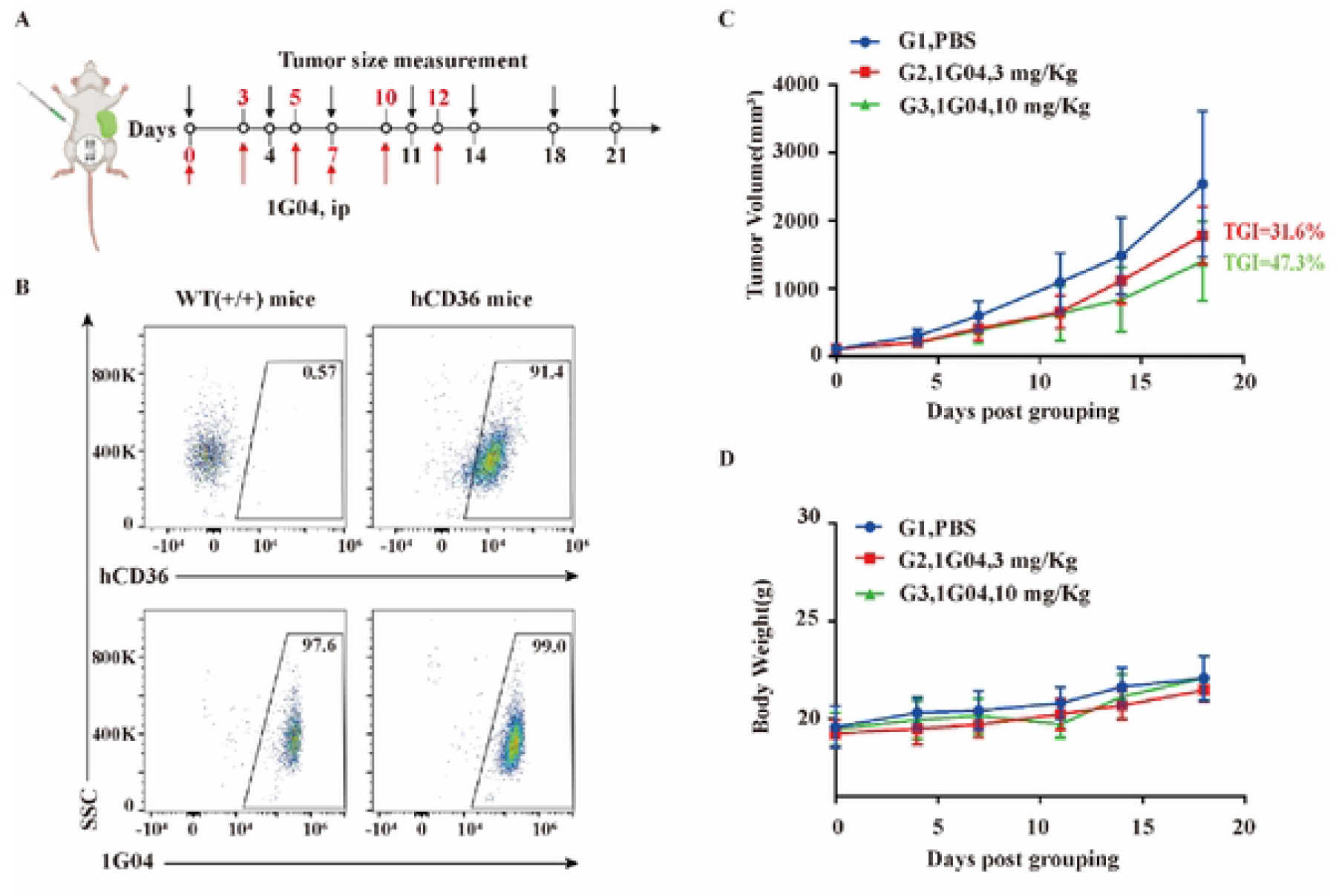
Validation of the efficacy of 1G04 in a hCD36 murine colon cancer model(n=3,6~8-week-old). A: Design of *in vivo* drug efficacy programs; B: Analysis of the *in vitro* binding of 1G04 to CD36. C: Tumor volume measurement during the treatment. D: Body weight record during the treatment.

## Discussion

CD36 is actively involved in the growth, tumor immunity, metastatic invasion, and drug resistance of various tumors through lipid metabolism[30]. Blocking CD36-mediated lipid metabolism is a strategy that can be considered for future tumor therapy. However, a lack of mouse models that faithfully recapitulate the human expression patterns of hCD36 has hindered progress on evaluating the anti-tumor effect of hCD36-targeting antibodies *in vivo*. Here, we developed a hCD36 mouse model by replacing the extracellular region of mCD36 with human ones, which allows testing the therapeutic potential of anti-hCD36-targeting antibody. Expression of hCD36 was only detectable in hCD36 mice while mCD36 was not expressed. Next, the distribution of lymphocyte subpopulations in the blood, lymph nodes and spleen of this mouse was consistent with that of wild-type mice, indicating that humanized mice did not affect the distribution of lymphocyte subpopulations. Furthermore, the immune function, blood cell composition and morphology, and vital organ function were not affected by humanization.

We further validated the inhibitory effect of 1G04 on colon cancer tumors in hCD36 mice. Our data show that targeting the hCD36 with blocking antibody is a promising strategy to enhance antitumor immunity and reduce tumor burden. However, the development of CD36 antibodies is mostly in the preclinical and biological testing stage, and there are few CD36 antibodies targeting tumor therapy. This is not only a limitation of the hCD36 mice in this study in the validation of CD36 tumor therapeutic anti-conductor efficacy but also a direction for us to continue to go deeper in the future. It is believed that with the emergence of drugs targeting metabolic inflammatory syndrome and the cardiovascular class of CD36 antibodies in future, hCD36 mice could also be able to further accelerate the development of this class of antibodies from preclinical to clinical progress. In addition, the use of humanized mice in human immuno-oncology research includes the exploration of mechanisms of action, preclinical pharmacodynamic and toxicological evaluation, and target discovery[19]. Thus, hCD36 mice could also further discover the potential association of CD36 with different targets in several diseases, thus facilitating drug development for these diseases.

## Conclusion

We have successfully generated hCD36 mice by a genetic engineering approach and validated the inhibitory effect of 1G04 on colon cancer tumor growth in these mice. We conclude that these hCD36 mice are a very useful tool for the preclinical evaluation of anti-CD36 therapeutic antibodies.

## Human and animal rights

The authors state that the procedures followed were according to the World Medical Association and the regulations of the responsible Clinical Research Ethics Committee.

## Authorship contributions

**Conceptualization:** Linlin Wang

**Formal analysis:** Linlin Wang and Zhenlan Niu

**Investigation:** Xingyan Yu

**Methodology:** Xiulong Xie

**Project administration:** Xiaofei Zhou and Yi Yang

**Supervision:** Hongyan Jing

**Writing – original draft:** Xiulong Xie

**Writing – review & editing:** Zhenlan Niu and Yi Yang

## Acknowledgements

The authors would like to thank staff of the Animal Center of Biocytogen Pharmaceuticals (Beijing) Co., Ltd. for breeding, housing and maintaining mouse.

## Supporting information

**S1 Fig. Expression of CD36 on immune cells in the blood of WT mice and hCD36 mice.**

**S2 Fig. Analysis of immune cells subpopulation in the blood of hCD36 and WT mice(n=3,8-week-old).** A, B: The frequency of T cells, B cells, NK cells, Monocyte, DC cell, and macrophage cells in mCD45 cells. C, D: The frequency of CD4+ T cells, CD8+ T cells, and Treg cells in mCD3 cells.

**S3 Fig. Analysis of immune cells subpopulation in the lymph nodes of hCD36 and WT mice(n=3,8-week-old)**. A, B: The frequency of T cells, B cells, NK cells in mCD45 cells. C, D: The frequency of CD4+ T cells, CD8+ T cells, and Treg cells in mCD3 cells.

